# RGS6 mediates exercise-induced recovery of hippocampal neurogenesis, learning, and memory in an Alzheimer’s mouse model

**DOI:** 10.1101/2023.04.17.537272

**Authors:** Mackenzie M. Spicer, Jianqi Yang, Daniel Fu, Alison N. DeVore, Marisol Lauffer, Nilufer S. Atasoy, Deniz Atasoy, Rory A. Fisher

## Abstract

Hippocampal neuronal loss causes cognitive dysfunction in Alzheimer’s disease (AD). Adult hippocampal neurogenesis (AHN) is reduced in AD patients. Exercise stimulates AHN in rodents and improves memory and slows cognitive decline in AD patients. However, the molecular pathways for exercise-induced AHN and improved cognition in AD are poorly understood. Here, we show that voluntary running in APP_SWE_ mice restores their hippocampal cognitive impairments to that of control mice. This cognitive rescue was abolished by RGS6 deletion in dentate gyrus (DG) neuronal progenitors (NPs), which also abolished running-mediated increases in AHN. AHN was reduced in sedentary APP_SWE_ mice versus control mice, with basal AHN reduced by RGS6 deletion in DG NPs. RGS6 expression is significantly lower in the DG of AD patients. Thus, RGS6 mediates exercise-induced rescue of impaired cognition and AHN in AD mice, identifying RGS6 in DG NPs as a potential target to combat hippocampal neuron loss in AD.

**Teaser:** RGS6 expression in hippocampal NPCs promotes voluntary running-induced neurogenesis and restored cognition in APP_SWE_ mice.

**Field Codes:** RGS6, Alzheimer’s disease, adult hippocampal neurogenesis, neural precursor cells, dentate gyrus, exercise, learning/memory

## Introduction

Alzheimer’s Disease (AD) is the most common neurodegenerative disorder and cause of dementia in older populations, accounting for 60-70% of dementia diagnoses(1). AD is a progressive neurological disease characterized by both cognitive and non-cognitive symptoms resulting from brain atrophy (2, 3). Hallmark neuropathological alterations in AD patients include accumulation of the protein fragment amyloid-beta (Aβ) outside of neurons and of hyperphosphorylated tau (p-tau) in neurofibrillary tangles within neurons. Accumulation of these pathological proteins causes massive neuronal loss prominently in the hippocampus, cortex, and other brain regions(4, 5). The hippocampus plays a critical role in learning and memory, and hippocampal degeneration is thought to underlie their progressive loss in AD. The only approved therapy for AD is lecanemab, an amyloid-targeting antibody that moderately reduces cognitive decline in early AD (6).

Despite the limited availability of neuroprotective interventions for AD, there is consistent evidence demonstrating beneficial effects of exercise on human AD pathogenesis and progression. Exercise protects against the risk of AD-induced dementia, improves memory, and slows cognitive decline in AD patients, as well as maintains hippocampal volume and prevents aging-related neurodegeneration (7–12). These effects of exercise are associated with increases in dentate gyrus blood flow that are correlated with postmortem measures of neurogenesis (13). Adult hippocampal neurogenesis (AHN) is a process occurring in the subgranular zone (SGZ) of the dentate gyrus (DG) that promotes the development and maturation of new granule neurons from neural stem cells and intermediate progenitor cells (IPCs) (14, 15). Numerous studies have demonstrated that exercise increases AHN in rodents (16–18).

After the discovery of AHN in rodents in 1963 (19), Eriksson (20) demonstrated its existence in humans by post-mortem analysis of brains from humans treated with the DNA incorporating drug BrdU. Spalding *et al.* (21) then demonstrated that AHN contributes to 700 new neurons/day in humans using carbon birth dating of hippocampal cells. Subsequent studies confirmed the existence of thousands of immature neurons in the dentate gyrus, but not other brain regions, of neurologically healthy humans up to the ninth decade of life by use of multiple cell markers, stereology, and other approaches (22–24). Recent single cell transcriptomic and immunocytochemical analyses identified all neurogenic populations in the DG of macaque, the exclusive existence of immature neurons in the DG of humans across the lifespan (25, 26), and the existence neurogenesis in *ex vivo* neurospheres from macaque and human DG (25, 26). Despite this evidence, there has been some controversy regarding the existence or extent of AHN in humans (27–29). It seems likely that the failure to detect AHN in some of these previous studies may be due to technical issues, including postmortem age, fixation, use of particular markers or epileptic tissue as has been discussed previously (22, 30, 31). While it is not possible to demonstrate causality between AHN and behavior in humans, the number of new neurons generated by AHN in humans is believed sufficient to impact the DG circuit (22). Thus, AHN may be a critical contributor in promoting hippocampal-dependent learning and memory and protect against cognitive loss in aging and AD(32–34). Indeed, reduced AHN has been linked to aging and neurodegeneration in both rodents and humans (14, 23, 32, 35-37), with AD patients exhibiting severely reduced AHN (23).

In mice, voluntary wheel running stimulates AHN and improves cognition (16, 17), an effect recently shown to be mediated by expression of RGS6 in DG NPCs (38). RGS6 is a member of the RGS protein family that negatively regulates Gα_i/o_-coupled GPCR signaling (39-43). Gao *et al* (38) showed that RGS6 promotes AHN and cognitive improvement in mice due to its negative regulation of GABA_B_Rs (40), whose activation would promote hyperpolarization and suppress AHN (14, 15). This is because GABA drives AHN by depolarizing neural progenitors and immature neurons in the brain (44). Though voluntary wheel running also improves cognition in mouse models of AD (45, 46), the extent to which running can modify AHN and cognition remains unknown. For example, though Choi *et al.*(45) found that running improved cognitive function in 5X FAD mice only after stratification of mice with the highest levels of AHN, the degree of improvement is unknown because mice without AD were not studied. Because fixed running wheels were not included for sedentary mice and the amount of running was not determined in either this study or that of Tapia-Rojas (46), it is also unclear whether differences were a result of running versus environmental enrichment. Further, Tapia-Rojas (46) found that even after running APP_SWE_/PS1ΔE9 mice showed significant cognitive impairment compared to control wild-type mice (46).

Here, we sought to determine the role of RGS6 in mediating voluntary running-induced AHN in the context of AD using Tg2576 (APP_SWE_) mice. We discovered that RGS6 is robustly expressed in the DG of humans and AD patients exhibit significant loss of RGS6-expression in the DG. Using retroviral approaches and novel RGS6-floxed mice, we generated RGS6^fl/fl^;APP_SWE_ mice to examine the impact of RGS6 deletion from DG neuronal progenitor cells (NPCs) on voluntary running-induced AHN and cognition in an AD mouse model. We show that voluntary running in APP_SWE_ mice completely restores impaired hippocampal learning and memory to that of control mice, and that this cognitive recovery is abolished by RGS6 deletion in DG NPCs. Finally, we demonstrate that RGS6 deletion from DG NPCs abolishes both basal and running-induced increases in AHN of APP_SWE_ mice. Given that exercise can improve or slow cognitive decline in AD patients and improve memory, maintain hippocampal volume, and slow aging-related degeneration, our findings identify RGS6 as a possible therapeutic target in AD and possibly aging related degeneration.

## Results

### Robust RGS6 expression in the dentate gyrus neurogenic niche of humans and its loss in AD patients

In our initial report of cloning transcripts encoding multiple splice forms of RGS6(47), we generated a peptide antibody specific for long N-terminal forms of RGS6 (RGS6L) to show robust expression of RGS6 in the mouse hippocampus, including the dentate gyrus (DG), the site of AHN. Given the recent finding that RGS6L mediates voluntary wheel running-induced AHN (38) and the link between exercise and cognitive improvements in AD patients (10, 12), we examined expression of RGS6L in the human DG. Fig. 1A shows that human DG expresses the same major RGS6 isoforms as seen in mice which are absent in the hippocampus of RGS6 global knockout mice. As we recently described (48), the lower 56kDa band represents RGS6L isoforms and the larger 69kDa band represents a novel brain-specific form of RGS6 (RGS6B) that contains the N-terminal sequence our antibody was generated against. There is a marked loss in RGS6L expression in the DG of AD patients (Figs. 1A-B). Immunohistochemical analysis further shows the expression of RGS6 in cells within the DG of humans and the significant loss of RGS6 expression in this region of the hippocampus of AD patients (Figs. 1C-D). These findings demonstrate that RGS6 is robustly expressed in the DG neurogenic niche, and that there is a significant reduction in RGS6 expression in the DG of AD patients. While the loss of RGS6 expression in the DG of AD patients may reflect hippocampal degeneration, its presence in the DG suggests it may play a key role in AHN in humans as it does in mice.

**Figure 1.**
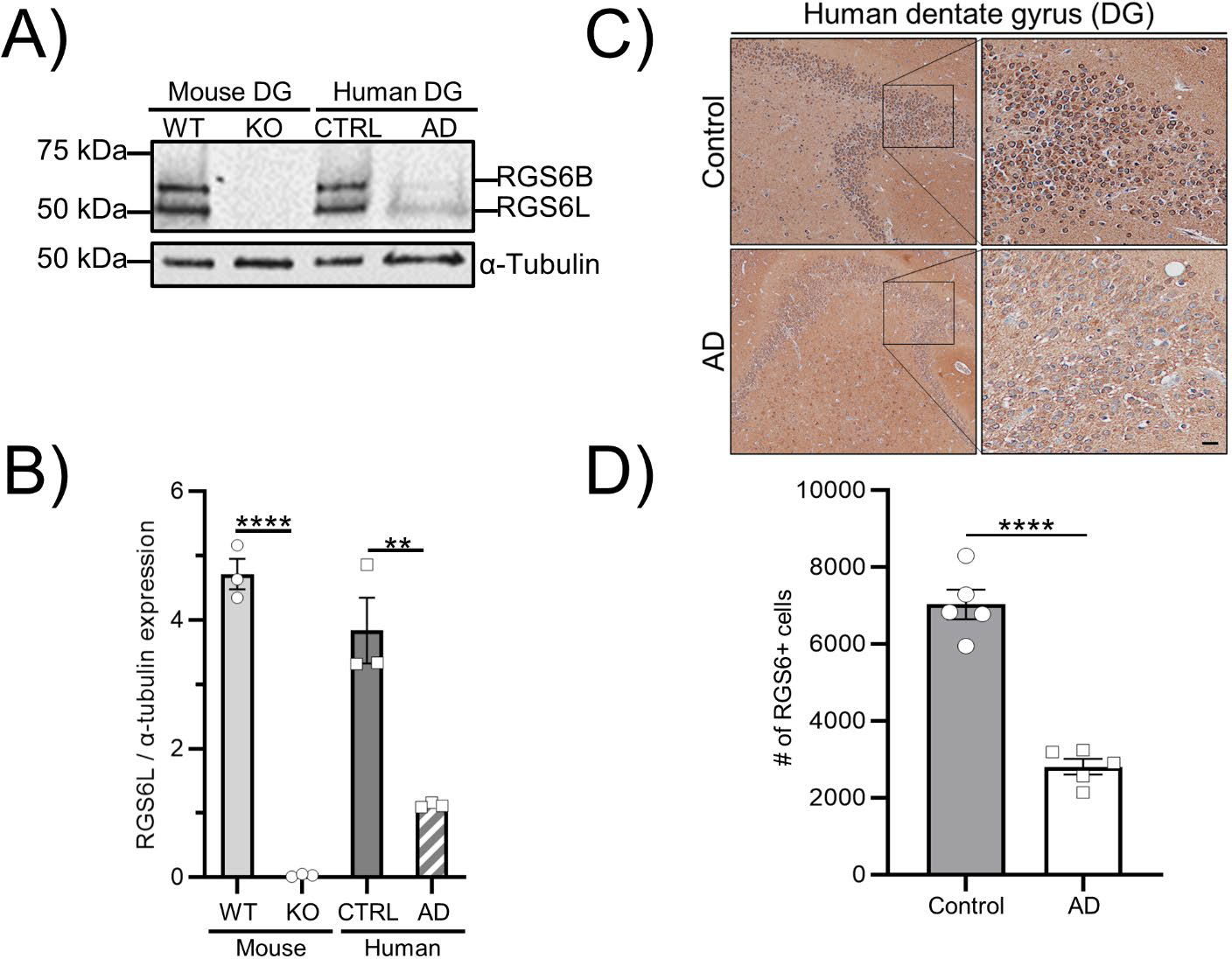
RGS6 is expressed in the human and mouse DG and is reduced in the DG of Alzheimer’s patients. **A-B)** Western blot (WB) analysis **(A)** and quantification **(B)** of RGS6L and α-Tubulin (loading control) co-expression in mouse (left lanes) and human (right lanes) dentate gyrus (DG) tissues. WT = wild-type, KO = global RGS6 KO, CTRL = control, AD = Alzheimer’s disease. One-way ANOVA was used to analyze mouse (light grey bars) and human (dark grey bars) separately. Significant effects of genotype (F_crit_ = 7.71, P = 0.0000) and diseases state (F_crit_ = 7.86, P = 0.006) were observed in mouse and human DG samples, respectively. **C-D)** Representative immunohistochemical (IHC) analysis **(C)** and quantification **(D)** of RGS6 expression in DG samples collected from age-matched neurotypical (control) and Alzheimer’s (AD) patients. Black boxes in **C** represent the region used to obtain the enlarged areas shown to the right. One-way ANOVA was used to analyze quantification data. A significant effect of disease was observed (F_crit_ = 5.318, P = 0.0000). *N* = 5 patients/group with an effect size of 6.13. ****P ≤ 0.0001, **P ≤ 0.01. Scale bar represents 1mm.

### RGS6 is required for exercise-induced cognitive improvements in an AD mouse model

Exercise, which stimulates AHN, has been shown to increase hippocampal volume and spatial memory in neurotypical elderly patients and significantly reduces the risk of AD development (8-12). Given that RGS6 in DG NPCs mediates voluntary running-induced AHN in mice (38), we next sought to determine whether exercise-mediated cognitive improvement in an AD mouse model was RGS6-dependent. To test this, we created RGS6^fl/fl^ mice (Fig. 2A) and crossed them with APP_SWE_ Tg2576 mice (Fig. 2B-C) to generate RGS6^fl/fl^, APP ^+/−^ mice (referred to as RGS6^fl/fl^, APP_SWE_).

**Figure 2.**
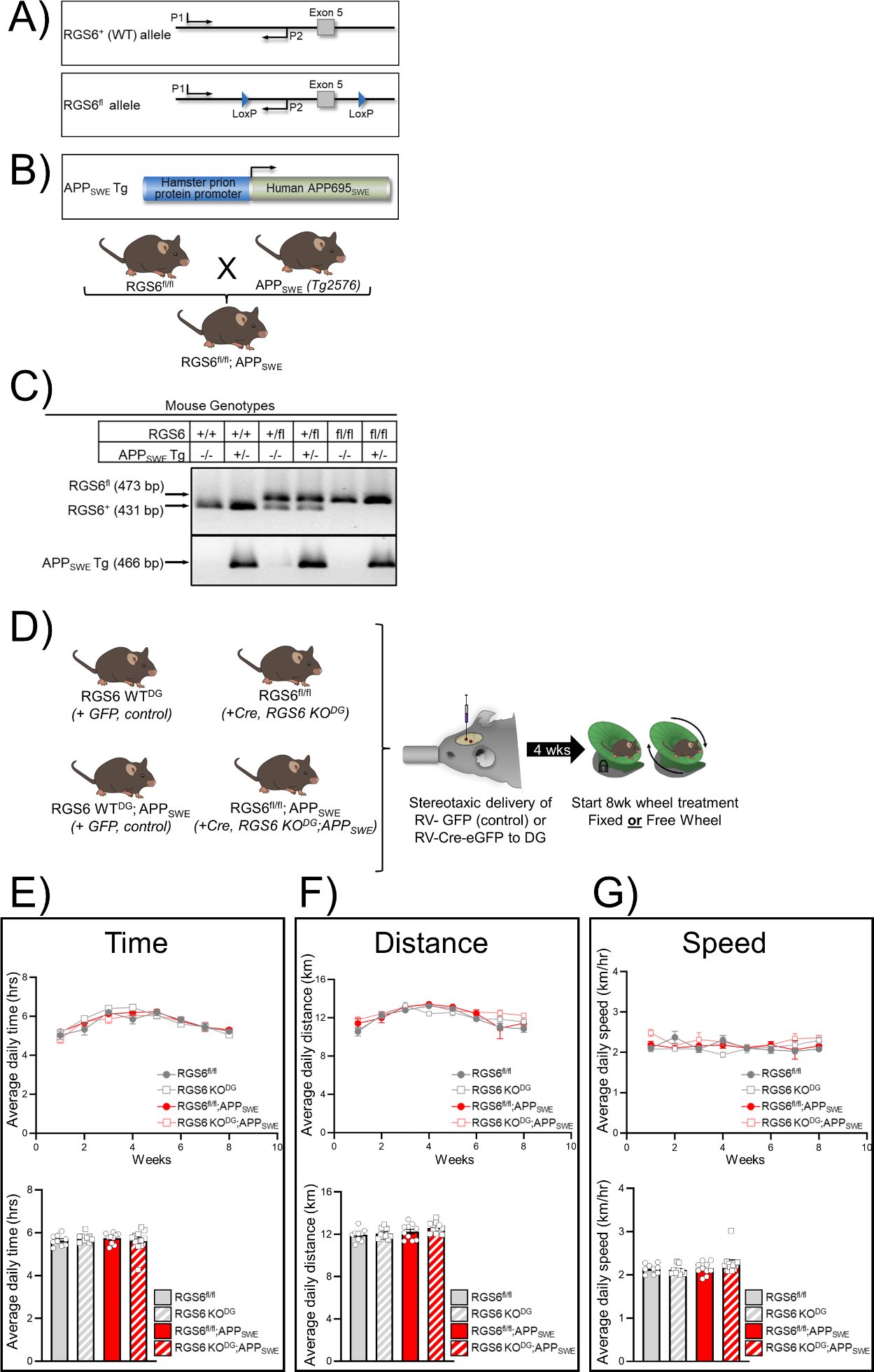
RGS6 loss in the DG does not impair the running patterns of mice. **A)** Schematic outlining CRISPR-Cas9 generation of RGS6^fl/fl^ mice. *Rgs6 WT* (*top*) and *Rgs6 flox* (*bottom*) alleles are depicted. E5, E6, E7 denote *Rgs6* exons 5-7. PCR primers 1 and 2 (arrows) were used to amplify a DNA segment in the *Rgs6* gene upstream of E5. **B)** Schematics outlining the APP_SWE_ transgene (*top*) and breeding cross to generate RGS6^fl/fl^;APP_SWE_ mice (*bottom*). **C)** Polymerase chain reaction (PCR) results confirming generation of RGS6^fl/fl^;APP_SWE_ mice. **D)** Schematic outlining running wheel set up, mouse genotypes, and running groups. **E-G)** Daily average running behavior patterns of 20 week old RGS6 WT^DG^ (dark grey closed circles/line), RGS6 KO^DG^ (light grey border squares/line), RGS6 WT^DG^;APP_SWE_ (red closed circles/line), and RGS6 KO^DG^;APP_SWE_ (light red border squares/line) mice on free running wheels over the course of 8wks during their dark cycle. Daily average time **(E)**, distance travelled **(F)**, and running speed **(G)** of mice during the dark cycle. Data are expressed as mean ± SEM. Two-way ANOVA with Tukey’s *post-hoc* adjustment was used to analyze the effects of and interaction between sex and genotype. No significant effects of genotype or sex were observed.

APP_SWE_ mice (Tg2576) are a transgenic AD mouse model that overexpress the Swedish mutation (KM670/671NL) of human APP695. This mutation causes early-onset AD in humans (49). APP_SWE_ mice exhibit behavioral and pathophysiological abnormalities recapitulating AD, including impaired spatial learning and memory, increased levels of Aβ42 and Aβ40, and amyloid deposits, as well as decreased hippocampal spine density, LTP, and volume (50, 51). APP_SWE_ mice also exhibit impaired AHN that is mediated by Aβ expression in neural stem cells at prodromal stages (1.5 mos.) (35, 52). Voluntary wheel running improves spatial learning and memory, decreases amyloid plaque formation, and increases hippocampal volume relative to sedentary APP_SWE_ mice (53–55).

To examine the role of RGS6 in DG NPCs on exercise-induced cognitive improvements in APP_SWE_ mice, we used stereotactic delivery of retrovirus (RV) expressing Cre to the SGZ of the DG of 3mo old RGS6^fl/fl^, APP_SWE_ mice or control RGS6^fl/fl^ mice. Because RV only infects proliferating cells, it is highly selective for NPCs when delivered to the DG SGZ, and therefore is used to deliver reporters, (*e.g.* eGFP) to measure AHN (14, 56-58). RV Cre delivery is also more selective than Cre mouse lines with promoters active in NPCs because these same promoters can show activity in other neurogenic as well as non-neurogenic brain regions (59). We used two injections into the DG that cover ∼ 70% of the DG SGZ region using the same stereotactic coordinates that were used to express dnWnt in the DG SGZ to abolish DG neurogenesis (60). Finally, we included eGFP in both the RV-Cre and RV-control to facilitate measurement of AHN in these mice by visualizing eGFP-positive neurons. We designated mice receiving eGFP and Cre-eGFP injections into the DG as RGS6 WT^DG^ and RGS6 KO^DG^, respectively (Fig. 2D).

Effects of voluntary wheel running on hippocampal-based learning and memory was performed in these four groups of mice by subdividing them into “fixed” (sedentary control) and “free” (running) wheel treatment groups for 8wks. Wheel running time, distance, and speed were recorded each day using Arduinos programmed to capture wheel turns. Daily averages of time (Fig. 2E), distance (Fig. 2F), and speed (Fig. 2G) were recorded in mice given free wheel access. Importantly, no significant effect of genotype was observed on the amount of time on wheel, distance travelled, or speed. Additionally, no significant effect of sex was observed on the running behavior of mice (Supplementary Fig. 2), therefore male and female running behavior data was combined (Fig. 2E-G). These data show that, if we observe any differences in learning and memory in mice on free running wheels, the differences cannot be attributed to differences in running behavior.

Working memory is one of the most well-modeled memory deficits in AD, and the most widely used paradigms in mice are maze-tasks which require spatial working memory to solve (61). The Y-maze spontaneous alternation test reports on spatial working or short-term memory. Hippocampal-dependent contextual fear conditioning (CFC) occurs when a context becomes associated with an aversive unconditioned stimulus like a foot shock (62). Nobili *et al.*(63) and others (51, 64) have demonstrated deficits in CFC in APP_SWE_ mice as a measure of hippocampal-based associative fear learning and memory.

We assessed hippocampal-based learning and memory in sedentary and running mice after 4 and 8 weeks of wheel access (Fig. 3A). Two-trial (2T) Y-maze spatial memory was tested at 4 and 8 weeks, and contextual fear conditioning (CFC) behavior was assessed at 8 weeks. Our results reveal a remarkable recovery of hippocampal learning and memory in APP_SWE_ mice to levels seen in control mice by voluntary wheel running that is completely dependent upon DG NPC RGS6 expression. As shown, voluntary wheel running significantly and dramatically improved learning and memory in APP_SWE_ mice without DG RGS6 deletion (RGS6 WT^DG^; APP_SWE_), as measured by spontaneous alternations and time spent in the novel arm during 2T Y-maze at both 4wks (Fig. 3B) and 8wks (Fig. 3C), as well as freezing behavior during CFC (Fig. 3D). While DG NPC-specific RGS6 loss alone impaired hippocampal-based learning and memory in control mice, we did not observe significant running-induced cognitive improvements in these mice. However, DG RGS6 loss in APP_SWE_ mice completely prevented running-induced improvements in the 2T Y-maze and CFC assessments. Interestingly, 8wks of running did not improve 2T Y-maze learning and memory in RGS6 WT^DG^; APP_SWE_ mice relative to 4wks (Supplementary Fig. 3), demonstrating that longevity of exercise over time does not significantly improve hippocampal-based spatial learning and memory. This is congruent with the running behavior data, as the mice do not run for significantly longer periods of time at 8 weeks versus 4 weeks (Fig. 2E). Previous literature has shown that running time, distance, and speed tends to plateau in male and female mice (C57BL/6J) after 3-4 weeks of running, consistent with our data (Fig. 2E-G).

**Figure 3.**
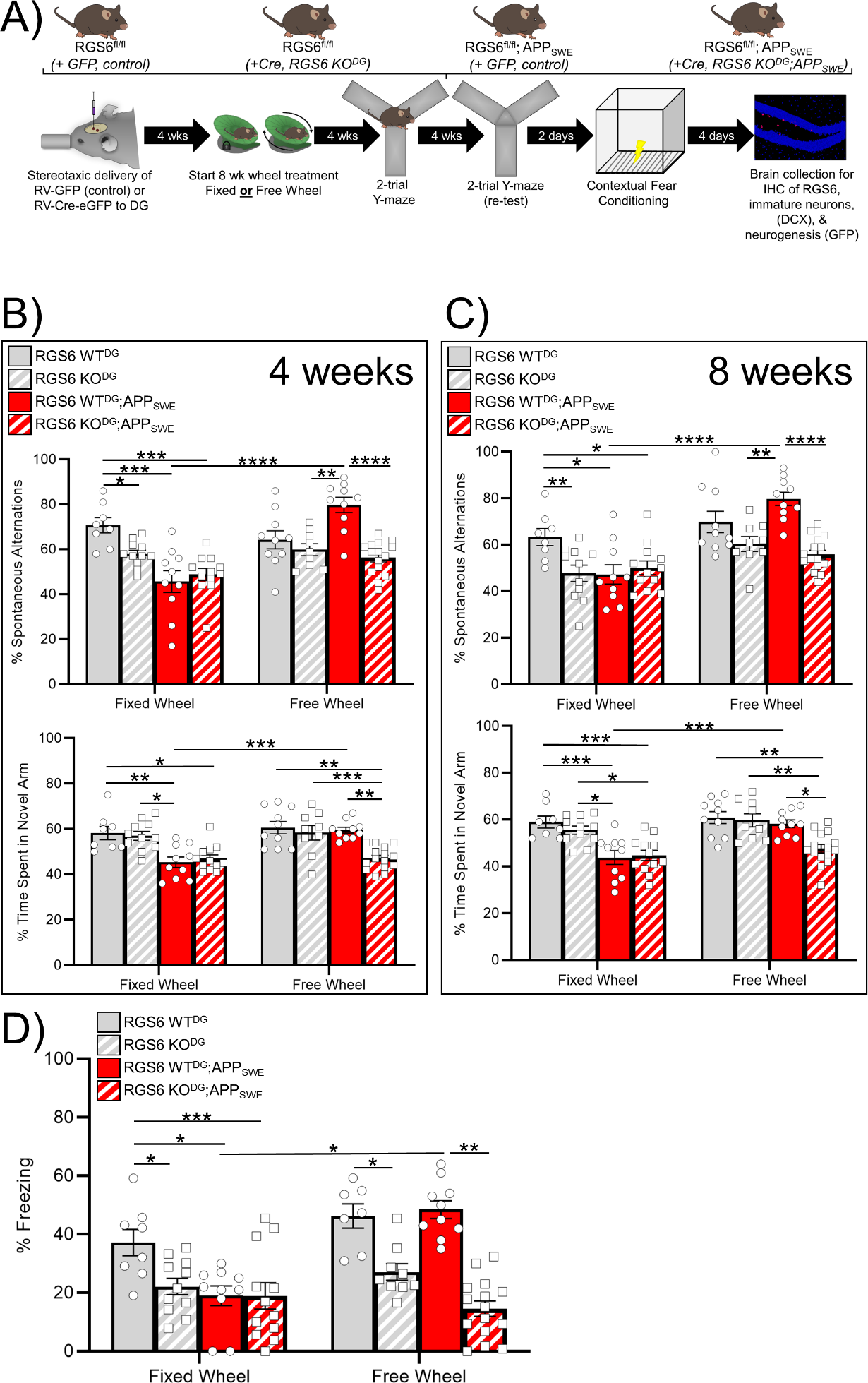
RGS6 in DG NPCs is required for exercise-induced cognitive improvements in APP_SWE_ mice. **A)** Schematic outlining experimental design. Mice were placed in cages containing either fixed (control) or free (running) wheels for 8 weeks. Mice underwent hippocampal-based learning and memory behavioral tasks after 4wks (Y-maze **(B)**) and 8wks (Y-maze **(C)** and contextual fear conditioning (CFC, **(D)**) of wheel treatment. **B-C)** Multi-way ANOVA with Tukey’s *post-hoc* adjustment was used to analyze the effects of and interactions between genotype, sex, time, and wheel treatment after 4 **(B)** or 8 weeks **(C)** of wheel treatment. <u>Top:</u> % Spontaneous alternations— Significant effects of genotype (*F_3_*_,154_ = 16.261, *P* = 0.000) and wheel treatment (*F_1_*_,154_ = 52.906, *P* = 0.000), as well as their interaction (*F_3_*_,154_ = 19.638, *P* = 0.000) were observed. No significant effects of sex or time were observed. <u>Bottom:</u> % Time spent in novel arm— Significant effects of genotype (*F_3_*_,154_ = 30.835, *P* = 0.000) and wheel treatment (*F_1_*_,154_ = 22.496, *P* = 0.000), as well as their interaction (*F_3_*_,154_ = 7.652, *P* = 0.000) were observed. No significant effects of sex or time were observed. **d** Multi-way ANOVA with Tukey’s *post-hoc* adjustment was used to analyze the effects of and interactions between genotype, sex, and wheel treatment. A significant effect of genotype (*F_3_*_,55_ = 6.217, *P* = 0.001) and its interaction with wheel treatment (*F_3_*_,55_ = 3.898, *P* = 0.013) were observed. No significant effects of sex or wheel treatment were observed. Data are expressed as mean ± SEM. \**P* ≤ 0.05, \*\**P* ≤ 0.01, \*\*\**P* ≤ 0.001.

### RGS6 is necessary for basal and running-induced AHN in APP_SWE_ mice

Given our behavioral evidence that RGS6 is required for running-induced reversal of cognitive dysfunction in APP_SWE_ mice, we evaluated the role of RGS6 in DG NPCs in AHN, which is reduced in AD in humans and amyloid mouse models of AD. We performed immunohistochemical analyses of the number of eGFP-(derived from RV-infected NPCs), doublecortin (DCX)- and neuronal nuclear antigen (NeuN)-positive neurons in hippocampal sections of the mice used for behavioral studies above (Fig. 4A). First, APP_SWE_ mice exhibited strong co-localization of eGFP and DCX immunoreactivity in neurons within the DG, identifying these as adult born neurons derived from DG NPCs (Fig. 4B). Second, voluntary running significantly increased the number of these adult born neurons and the number of NeuN positive neurons in RGS6 WT^DG^; APP_SWE_ mice (Fig. 4B-C). Third, RGS6 deletion from DG NPCs significantly impaired AHN in both sedentary and running RGS6 WT^DG^ and RGS6 WT^DG^; APP_SWE_ mice (Fig. 4B-C; Supplementary Fig. 4). Finally, while control RGS6 WT^DG^ mice had higher basal AHN relative to their APP_SWE_ counterparts, running did not significantly increase AHN in these mice (Fig. 4B-C; Supplementary Fig. 4). Again, RGS6 deletion significantly impaired AHN in these mice as in their APP_SWE_ counterparts. These results show that running-induced cognitive improvements in APP_SWE_ mice are accompanied by increases in AHN and adult born neurons that are dependent upon RGS6 expression in DG NPCs. There was a similar correlation between AHN and learning and memory in RGS6 WT^DG^ mice where we observed no effects of running on cognition or AHN but where RGS6 deletion from DG NPCs caused loss of both cognition and AHN (Fig. 3B-D; Fig. 4B-C; Supplementary Fig. 4). These findings show that RGS6 is necessary for basal and voluntary running-induced AHN in APP_SWE_ mice.

**Figure 4.**
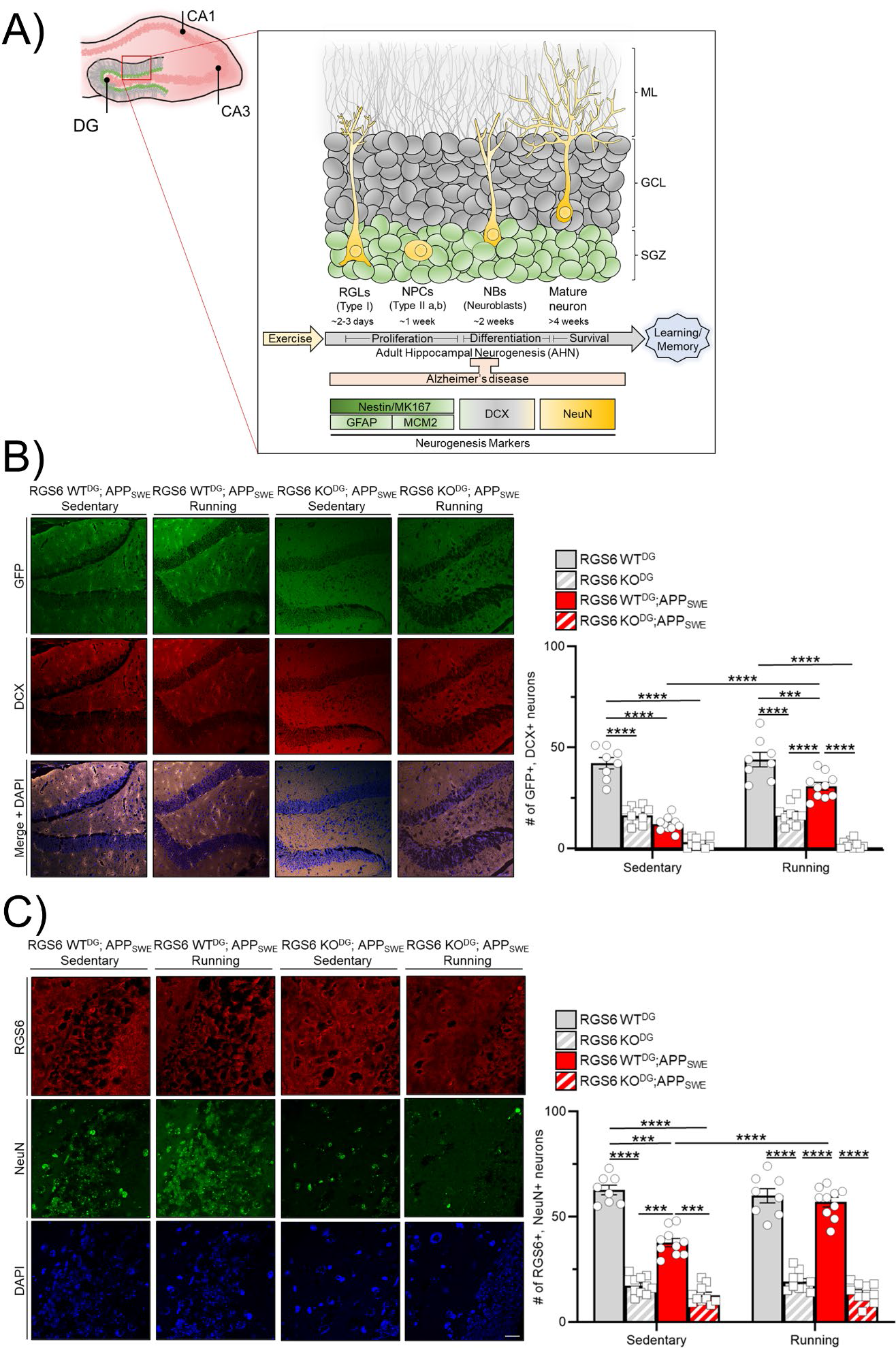
RGS6 is necessary for basal and running-induced AHN in APP_SWE_ mice. **A)** Schematic summarizing the process of AHN (*top*) and markers used at each stage (*bottom*). **B)** <u>Left:</u> Representative IF images (20x, *N* = 10/group) of neurogenesis markers (RV-eGFP or RV-Cre-eGFP, green; doublecortin (DCX), red) in RGS6 WT^DG^; APP_SWE_ and RGS6 KO^DG^; APP_SWE_ sedentary and running mice from Figure 3. Scale bar is 25µm. <u>Right:</u> Quantification of the number of GFP+ DCX+ neurons in RGS6 WT^DG^, RGS6 KO^DG^, RGS6 WT^DG^;APP_SWE_, and RGS6 KO^DG^; APP_SWE_ mice. Multi-way ANOVA with Tukey’s *post-hoc* adjustment was used to analyze the effects of and interactions between genotype, wheel treatment, and sex. Significant effects of genotype (*F*_3,68_ = 157.885, *P* = 0.000), wheel treatment (*F*_1,68_ = 14.969, *P* = 0.000), and their interaction (*F*_3,68_ = 13.633, *P* = 0.000) were observed. **C)** <u>Left:</u> Representative IF images (40x, *N* = 10/group) analyzing RGS6 (red) co-expression with NeuN (green) and DAPI (blue) in RGS6 WT^DG^; APP_SWE_ and RGS6 KO^DG^; APP_SWE_ sedentary and running mice from Figure 3. Scale bar is 40µm. <u>Right:</u> Quantification of the number of RGS6+ NeuN+ neurons in RGS6 WT^DG^, RGS6 KO^DG^, RGS6 WT^DG^;APP_SWE_, and RGS6 KO^DG^; APP_SWE_ mice. Multi-way ANOVA with Tukey’s *post-hoc* adjustment was used to analyze the effects of and interactions between genotype, wheel treatment, and sex. Significant effects of genotype (*F*_3,68_ = 264.564, *P* = 0.000), wheel treatment (*F*_1,68_ = 12.290, *P* = 0.001), and their interaction (*F*_3,68_ = 12.621, *P* = 0.000) were observed. Data are expressed as mean ± SEM. \*\*\**P* ≤ 0.001, \*\*\*\**P* ≤ 0.0001.

## Discussion

AD is characterized by brain atrophy, including massive neuronal loss in the hippocampus, which is responsible for the learning and memory deficits associated with this disease. AHN is severely reduced in AD (23) which may be an important contributor to hippocampal atrophy in AD. In contrast, exercise, a known stimulator of AHN, produces significant improvements in cognitive function in AD patients and in AD mouse models. Thus, there is a critical need to understand the AHN mechanisms that contribute to improved hippocampal function in AD. Here, we have addressed a gap in our understanding of how exercise may improve cognitive function in AD by showing that voluntary running-induced AHN and reversal of impaired cognitive function in an AD mouse model is mediated by RGS6 in DG NPCs.

Our results demonstrate that RGS6 in DG NPCs mediates the ability of voluntary running to completely restore impaired hippocampal learning and memory in a mouse model of AD. The restoration of cognitive dysfunction in APP_SWE_ mice by voluntary running was not due to differences in running patterns or sex. Moreover, we found that 8wks of running did not improve learning and memory in mice relative to 4wks, demonstrating that longevity of exercise over time does not significantly improve hippocampal-based spatial learning and memory. Our finding that DG NPC RGS6 loss abolishes running-induced AHN in these AD mice suggests that RGS6 may combat hippocampal neuron loss and resultant cognitive dysfunction by promoting AHN.

AHN can be stimulated by exercise and environmental enrichment, while reduced AHN has been linked to aging and neurodegeneration in both rodents and humans (14, 23, 32, 35-37). Moreno-Jiminez (23) showed that AHN drops sharply in patients with AD. In mice, voluntary wheel running stimulates AHN and improves cognition (16, 17). Voluntary running also has these effects in mouse models of AD (45, 46). However, these studies have not revealed the extent of cognitive dysfunction within their mouse models of AD nor whether exercise can restore normal cognitive function due to lack of inclusion of non-AD mice and proper controls in these studies. Two studies have reported that voluntary running in AD mice improves spatial learning/memory to that of control mice (53, 54) comparing latency times in radial arm water maze (RAWM) or Barnes maze. However, neither study showed this was due to running by including locked running wheels in their studies, a complicating issue because one of these groups previously showed that AD mice with locked running wheels showed the same improvements in RAWM performance as those with free wheels-demonstrating this was not a running-mediated effect (65). The most definitive study by Choi *et al*.(45) demonstrated that 5XFAD mice exhibited increased AHN, reduced Aβ pathology, and modest improvement in spatial learning when stratified for running mice who showed AHN. That is, a subset of running mice not exhibiting AHN had no cognitive improvement. However, the extent of both cognitive impairment in these 5XFAD mice and cognitive rescue by exercise in this study is not known as no comparisons to wild type mice were made. In addition, sedentary mice were exposed to an environment lacking running wheels vs fixed running wheels, making it difficult to know if the effects on cognition were due to an enriched environment or running, which was not measured.

Our results reveal that RGS6 plays a critical role in both basal and running-induced AHN in mice. First, we found that RGS6 WT^DG^; APP_SWE_ mice had significantly reduced AHN compared to RGS6 WT^DG^ mice (Fig 4; Supplementary Fig 4). Thus, these mice display reduced AHN as observed in human AD patients. Second, RGS6 deletion from DG NPCs completely prevented running-induced increases in adult born neurons in APP_SWE_ mice, demonstrating that RGS6 is required for running-induced AHN. Third, though we did not observe increases in AHN in response to running in RGS6 WT^DG^ mice, RGS6 deletion from DG NPCs in these mice significantly reduced adult born neurons in both sedentary and running mice. The selectivity of the effect of voluntary running on increasing AHN in APP_SWE_ mice suggests that our conditions of voluntary running rescued the impaired AHN in these mice but that the level of RGS6 and ongoing AHN in RGS6 WT^DG^ mice is sufficient and not further increased by these conditions. This would suggest that diminished AHN in APP_SWE_ mice is more sensitive to running-induced increases in AHN or alternatively, that our running paradigm has a ceiling effect in terms of increasing AHN. Though Gao *et al*. (38) observed running-induced increases in AHN and cognition in wild-type mice, these analyses were performed following voluntary running for 17-18 days and 27 days, respectively, and the running activity of these mice was not measured as in the present study. Fourth, the lower level of AHN in APP_SWE_ mice may reflect a reduction in the number of RGS6-expressing NPCs in these mice, something that future studies could address by RNA seq. Finally, we observed a strong correlation between effects of running as well as RGS6 deletion from DG NPCs on AHN. The reversal of impaired hippocampal learning and memory in RGS6 WT^DG^; APP_SWE_ mice and suppression of it by RGS6 deletion from DG NPCs correlated well to changes in AHN under these conditions (Fig. 3-4; Supplementary Fig. 4). Likewise, the lack of running effects on hippocampal cognition in RGS6 WT^DG^ mice was accompanied by a lack of effects on AHN, whereas deletion of RGS6 from DG NPCs in these mice reduced both hippocampal-based learning and memory and AHN.

RGS6 is likely promoting cognitive improvements in APP_SWE_ mice by mediating AHN, though future studies are required to directly test this hypothesis. As an RGS protein, RGS6 functions as an essential negative modulator of GPCR signaling due to its ability to inactivate Gα_i/o_ protein signaling via its GTPase-activating activity towards the Gα subunit. RGS6 loss, which normally negatively regulates Gα_i/o_ signaling, leads to enhanced Gα_i/o_ signaling. This is important in the context of AHN as GABAergic signaling is important for neuronal precursor proliferation, differentiation, and migration, as well as their integration into synapses (66). In this process GABA switches to a depolarizing stimulus, so RGS6, by suppressing GABA_B_R neuronal silencing, would facilitate this process (15, 44). Gao *et al.* *(38)* showed that both voluntary running and increased RGS6 expression in NPCs reduced GABA_B_R-mediated suppression of Ca^2+^ currents, suggesting this would provide strong synergism with GABA_A_R-mediated depolarization involved in AHN. Taken together, we speculate that RGS6 may be promoting running-induced AHN by negatively regulating GABA_B_ inhibitory signaling to subsequently improve cognition in APP_SWE_ mice.

Overall, our results have led us to propose that RGS6 promotes voluntary running-induced cognitive improvements in APP_SWE_ mice by mediating AHN (Fig. 5). Given that exercise can protect against the risk of dementia due to AD, to improve or slow cognitive decline in AD patients, to maintain hippocampal volume, to improve memory and to delay or prevent aging-related neurodegeneration (7–12) our findings identify RGS6 as a possible therapeutic target in AD and possibly aging related degeneration.

**Figure 5.**
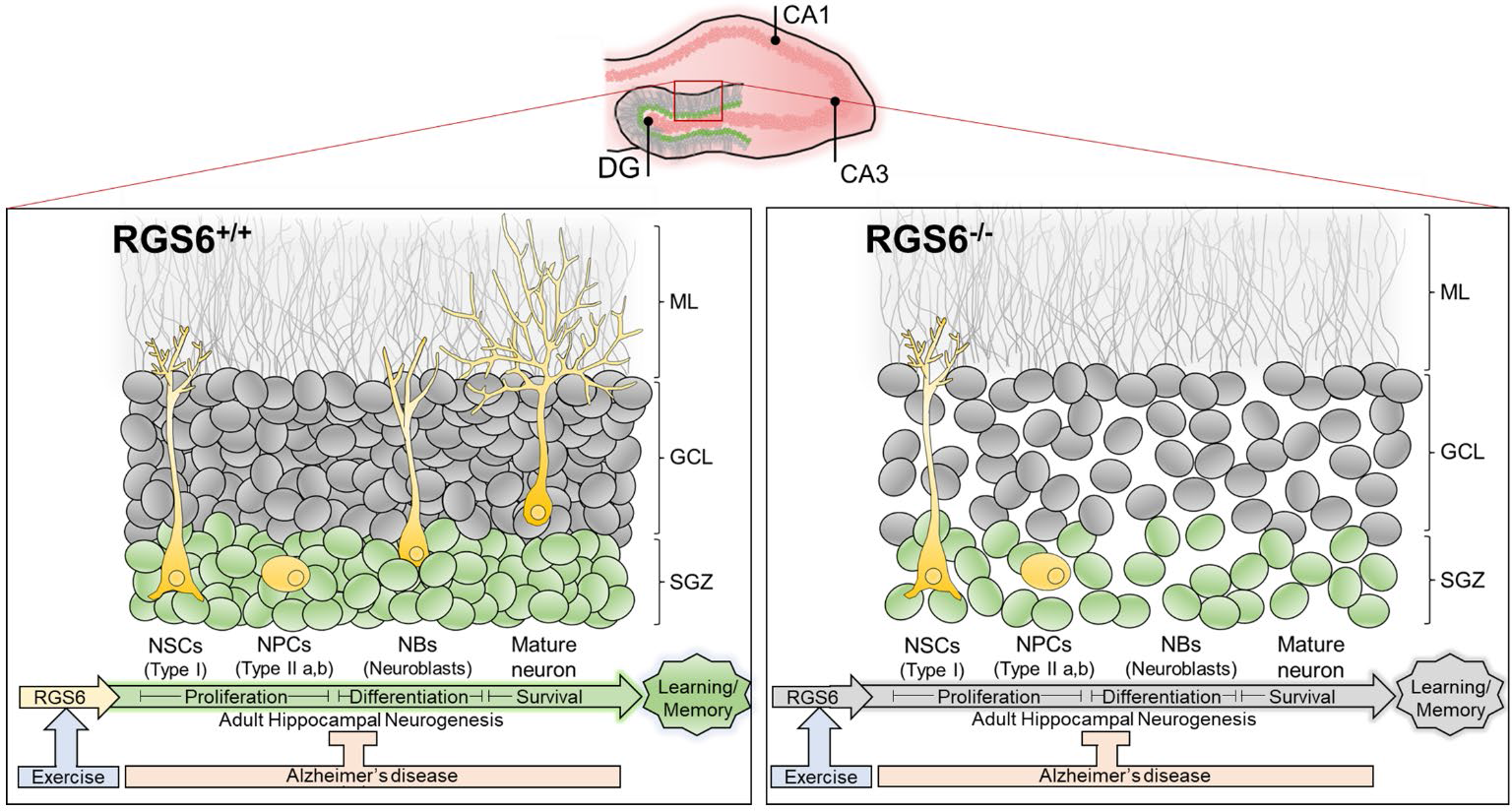
Summary model of RGS6’s role in DG NPCs. <u>Left:</u> RGS6 DG NPC expression, stimulated by voluntary running exercise, promotes AHN to improve cognitive function in AD mice. <u>Right:</u> RGS6 DG NPC loss leads to significant impairments in AHN, thus preventing the ability of voluntary running-mediated AHN to reverse cognitive dysfunction in AD mice.

## Acknowledgements

We would like to acknowledge Mariah Leidinger at the University of Iowa Pathology Core Facility and Dr. Li-Chun (Queena) Lin at the University of Iowa NeuroBank Core Facility for their contributions to the preparation of the human patient DG samples. All data needed to evaluate the conclusions in the paper are present in the paper and/or the Supplementary Materials. The data can be provided by the corresponding author pending scientific review and a completed material transfer agreement. Requests for the data should be submitted to. This research was supported by the National Institutes of Health (#AA025919, #AA025919-03S1, #AA025919-05S1).

## Inclusion and diversity

We support inclusive, diverse, and equitable conduct of research. One or more of the authors of this paper self-identifies as an underrepresented ethnic minority in their field of research or within their geographical location. One or more of the authors of this paper self-identifies as a gender minority in their field of research. One or more of the authors of this paper self-identifies as living with a disability.

## Materials and Methods

### Mice

All studies employed RGS6 floxed (RGS6^fl/fl^) and RGS6^fl/fl^; APP_SWE_ mice with and without retroviral-mediated RGS6 deletion in DG NPCs. RGS6^fl/fl^ mice were created on a mixed background (C57BL/6J x SJL) using CRISPR-Cas9 gene editing to target exon 5 for deletion of all known RGS6 splice forms(47). Mice were then backcrossed at least 5 generations to establish a C57BL6/J background. RGS6^fl/fl^ mice were bred with APP_SWE_ mice (Taconic 1349, C57BL/6J background) to create mice to determine the impact of RGS6 loss in the dentate gyrus (DG) on neurogenesis in AD mice. Mice used in voluntary wheel running studies (20wks) were housed in a room maintained at 22µC and 20-30% humidity on a reverse light cycle (12hrs light: 12hrs dark-lights off at 8:00am) and were given *ad libitum* access to food and water. All *in vivo* experiments were approved and performed in accordance with guidelines set by the Institutional Animal Care and Use Committee (IACUC) at the University of Iowa.

### Exercise-induced neurogenesis and behavioral studies

For running wheel studies, mice were housed individually and acclimated to wheels present in their cages 1 week prior to starting the running behavior. Cages contained either fixed or free running wheels (plastic, 5 × 5 × 3.5 inches, Chewy Inc.) affixed to the cage bottom (Supplementary Fig. 1). A single round hole magnet (10×3mm) facing outwards was glued to the side of each wheel. Electrical tape was used to affix an Arduino magnetic detector to the side of each cage, less than 2cm from the wheel magnet. Each magnetic detector was equipped with 3 jumper cables (5V, ground, digital output) that were coupled to an Arduino MEGA board (digital output) and breadboard (5V and ground). The Arduino MEGA board was coupled to a laptop equipped with Arduino IDE and PuTTY open-source software, and recorded running wheel data every 10 minutes for 8 consecutive weeks. Data and code sharing are available upon request.

### Hippocampal-dependent learning and memory behavioral tests

All mice were handled for 5 consecutive days in the respective behavior room prior to beginning training/testing. Mice were acclimated to the room 1hr prior to beginning training/testing trials. All behavior was conducted during the dark cycle and behavioral testing rooms remained dark as to not disrupt the dark cycle. The experimenter was blinded, and all mouse groups were randomized. Male and female mice were used for all studies.

#### Two-trial Y maze

The two-trial Y maze consists of a training trial followed by a testing trial (ITI = 2hrs) (Fig. 3). At the start of each trial, mice are placed in the designated “starting arm” of the maze, facing the back wall. During training, one arm of the Y maze was blocked off. Mice were allowed to freely explore the remaining two arms for 10min. Mice were tested 2hrs after completing their training trial. During testing, all arms of the Y maze are open, and mice were allowed to freely explore the maze for 5min. The number of arm entries and time spent in the previously blocked arm were recorded, as well as the number of spontaneous alternations.

#### Contextual Fear Conditioning (CFC)

CFC consists of three total trial phases: training, test 1, and test 2 (Supplementary Fig. 3C). For each trial, mice are placed in a fear conditioning box (Med Associates NIR VFC System) with a steel wire bottom. The training phase is 3min long with a 1.5mA electrical foot shock delivered at 2:30. The percentage of freezing behavior is recorded for 30sec post-shock and is considered the unconditioned/baseline fear behavior. The percent freezing during the 30sec immediately following shock is compared to the percent freezing behavior prior to shock delivery to ensure mice correctly respond to the shock. 24hrs following training, mice are placed in the same fear box and freezing behavior is recorded for 5min (Test 1), where no shock is delivered during the trial. Percent freezing during Test 1 is compared across all mouse groups to assess learning and memory of the shock treatment delivered during training. For Test 2, the walls and floors of the fear boxes are changed, and mice are placed in the fear boxes 24hrs following Test 1, and freezing behavior is recorded for 5min, where no shock is delivered. Percent freezing behavior during Test 2 is compared to freezing behavior during Test 1 to ensure the freezing behavior during Test 1 is context-specific, as the changed context during Test 2 should reduce freezing behavior from Test 1 (but will not reduce it to pre-shock freezing behavior).

### Stereotaxic Injections

Anesthesia (isoflurane) was induced at 5% and maintained at 2.5% throughout the surgery. Mice were oriented in the stereotaxic frame (Leica Angle One) and ear bars were placed to level and secure the head. Given that the DG is a large structure, mice for running wheel studies underwent craniotomies at 2 separate sites using stereotaxic coordinates (Site 1: AP = −2.06mm, ML = +/−1.25mm, DV = −1.92mm; Site 2: AP = −3.16mm, ML = +/−2.25mm, DV = −2.63mm; Franklin & Paxinos Mouse Brain Atlas), resulting in 4 total sites per mouse. Viral injections (1µL, Hamilton pipette) of either retrovirus (RV)-eGFP (control) or RV-Cre-eGFP (were delivered bilaterally at an infusion rate of 0.1µL/min. Mice received Buprenorphine-SR (2 mg/kg, i.p.), and head staples were used to close the surgical site for 2wks following surgery.

Mice continued to receive analgesic treatment once daily for 3 days following surgery and were monitored closely for 1 week following surgery. RV-eGFP and RV-Cre-eGFP were purchased from the Salk Institute GT3 Viral Core.

### Immunohistochemistry (IHC)

IHC analysis of RGS6 expression in human DG tissue. The University of Iowa Neurobank Core Facility provided 5 neurotypical human control DG tissues and 5 age-matched human AD patient DG tissues. Tissues were formalin-fixed paraffin-embedded (FFPE) and deparaffinized with a series of xylene and EtOH washes prior to incubation. Sections were incubated overnight at 4µC with rabbit-anti-RGS6 (homemade polyclonal antibody against the entire RGS6 protein (41, 42, 48), 1:200. Samples were imaged using an Olympus VS200 Slide Scanner (brightfield). Images were quantified using the FIJI “Cell Counter” analysis Plugin. *Post-hoc* power analyses (G*Power) indicate our effect size (6.13) for *N* = 5 patients/group produces a power of 99% at α =0.05.

### Immunofluorescence (IF)

IF analysis of RGS6 expression and neurogenesis markers was performed in mouse DG tissue samples. In preparation for IF, mouse tissues mice were perfused as previously described (39). Briefly, mice were first euthanized with isoflurane and immediately perfused (cardiac) with ice-cold 1X PBS followed by 4% paraformaldehyde (PFA). Brain tissues were removed, postfixed in 4% PFA at 4µC overnight, and finally sunk in a 30% sucrose solution (in 1X PBS). 10 µm coronal DG sections were collected (Olympus cryostat) and mounted. Tissue sections were blocked for 1hr at 4µC in 1X PBS + 10% goat serum + 0.5% Triton X-100 (blocking solution). After blocking, tissue sections were incubated overnight at 4µC in rabbit-anti-RGS6 (homemade polyclonal antibody against the entire RGS6 protein(41, 42, 48), 1:100, mouse-anti-GFP (Invitrogen, MA5-15256, 1:250), mouse-anti-DCX (Santa Cruz Sc-271390, 1:200), or mouse-anti-NeuN (Millipore MAB377, 1:200) primary antibody diluted in the blocking solution. Following overnight incubation, tissues were washed 5 times at room temperature in 1X PBS and subsequently incubated for 1hr at 4µC in Alexa Fluor 488/568/647-conjugated secondary fluorescent antibodies. Slices were mounted with ProLong Diamond Antifade Mountant with DAPI (Fisher Scientific) and imaged using an Olympus FV3000 confocal microscope. Images were quantified using the FIJI “Cell Counter” analysis Plugin.

### Western blotting (WB)

WB analysis was performed as we previously described(39). Frozen (−80µC) dentate gyrus samples were placed in a RIPA lysis buffer (150 mM NaCl, 1.0% NP-40, 0.5% sodium deoxycholate, 0.1% SDS, 50 mM Tris-HCL [pH 8.0] containing protease and phosphatase inhibitors (MilliporeSigma, P8340/P5726). Tissues were homogenized using a Dounce homogenizer and centrifuged at 14,000 g for 10 minutes (4°C) to isolate the protein-rich supernatant. The supernatant was then combined with Laemmli buffer, boiled for 5 minutes (95µC), loaded onto a 4% to 20% SDS/PAGE gel (BIO-RAD #4568094), and transferred to a nitrocellulose membrane. Following transfer, membranes were probed with rabbit anti-RGS6 (homemade polyclonal antibody against the entire RGS6 protein; 1:2500) and mouse anti–α-Tubulin (1:10,000; MilliporeSigma, CP06) primary antibodies, and Alexa Fluor 800–conjugated goat anti-rabbit (LI-COR Biosciences 680RD) and goat anti-mouse (LI-COR Biosciences 800CW) secondary antibodies. WBs were visualized using the LI-COR Odyssey 9120 Imaging System.

### Statistical Analyses

All data are expressed as mean ± SEM. One-way ANOVA was used to analyze IHC and IF quantification data (Fig. 1; Fig. 5). Two-way ANOVA with Tukey’s *post-hoc* adjustment was used to analyze the effects of and interaction between genotype and sex (Fig. 2; Supplementary Fig. 2). Multi-way ANOVAs were used to analyze the effects of and interactions between genotype, wheel treatment, and sex (Fig. 3D; Fig. 4; Supplementary Fig. 4), or genotype, wheel treatment, sex, and time (Fig. 3B-C; Supplementary Fig. 3). Sex as a biological variable was accounted for in all statistical analyses. Animal cohort sizes were determined from preliminary studies and G*Power analyses. *P* ≤ 0.05 was considered statistically significant. Statistical analyses were performed using XLSTAT software, and graphs were created using GraphPad Prism.

---

**Supplemental Figure 1.**
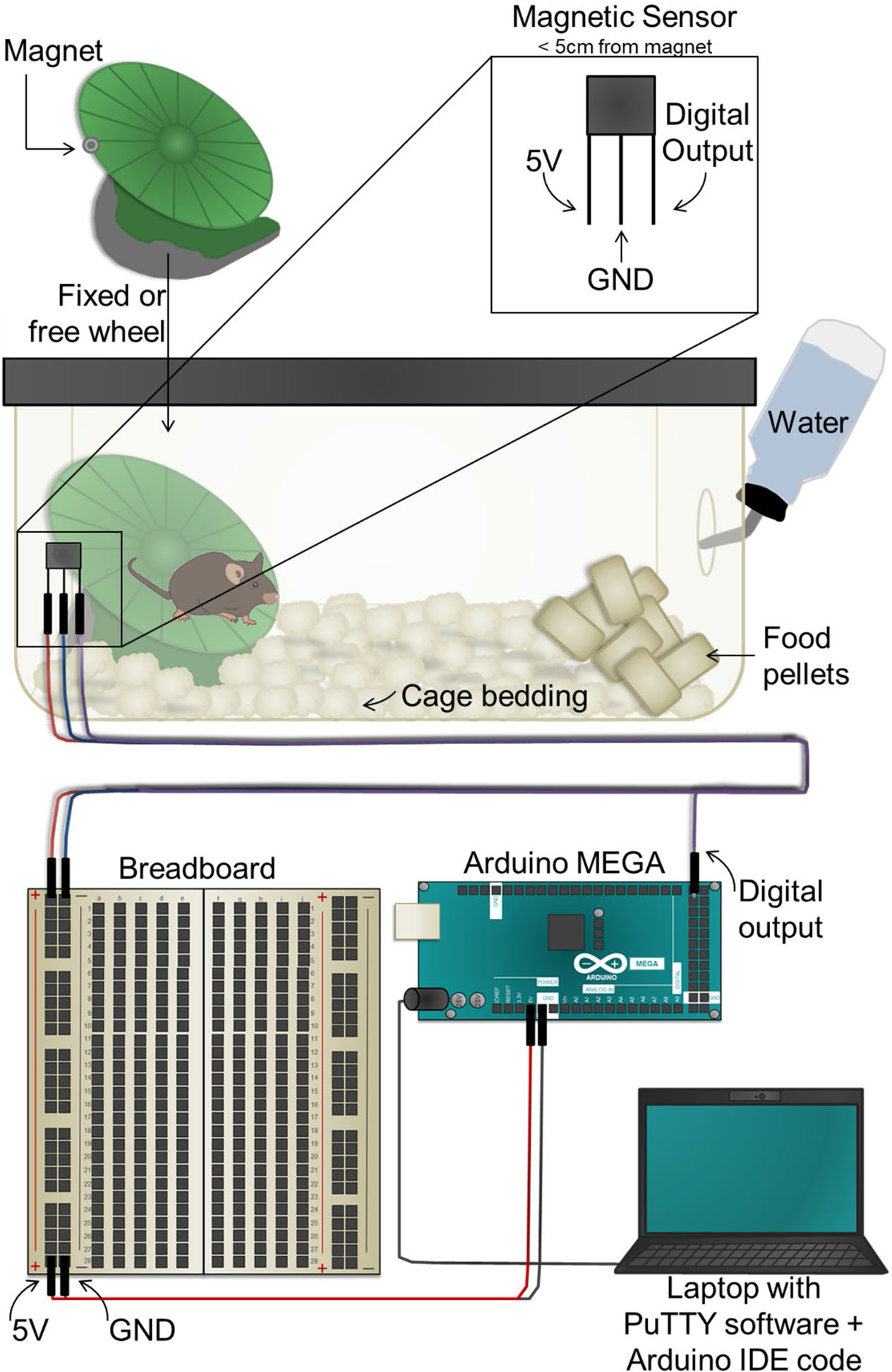
(Related to Fig. 2-3) *Arduino-based set up to record running pattern in mice*. Schematic detailing the cage, wheel, wiring, and Arduino set ups for the experiments performed in Figures 2 and 3. Mice were individually housed, with each cage containing either a free or fixed wheel affixed to the cage bottom via temporary glue. A single round hole magnet (10×3mm) facing outwards was glued to the side of each wheel. Electrical tape was used to affix an Arduino magnetic detector to the side of each cage, less than 2cm from the wheel magnet. Each magnetic detector was equipped with 3 jumper cables (5V, ground, digital output) that were coupled to an Arduino MEGA board (digital output) and breadboard (5V and ground). The Arduino MEGA board was coupled to a laptop equipped with Arduino IDE and PuTTY open-source software, and recorded running wheel data every 10 minutes for 8 consecutive weeks.

**Supplemental Figure 2.**
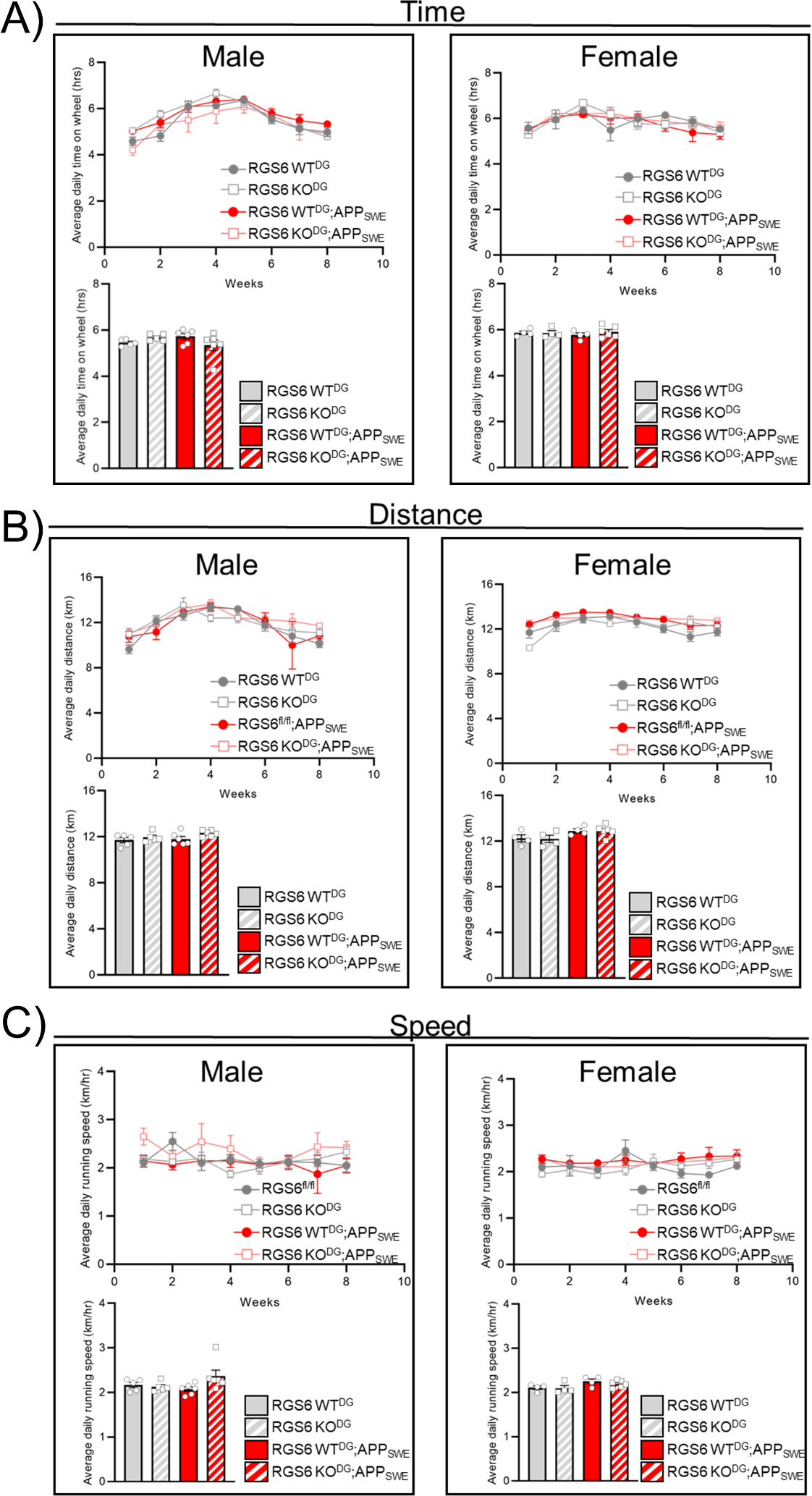
(Related to Fig. 2) *RGS6 loss in the DG does not impair the running patterns of mice*. **A-C)** Running patterns in male (*left panels*) and female (*right panels*) mice on free running wheels over the course of 8wks. Average time (A), distance (B), and speed (C) were recorded. Data are expressed as mean ± SEM. Two-way ANOVA was used to analyze the effects of and interaction between genotype and sex.

**Supplemental Figure 3.**
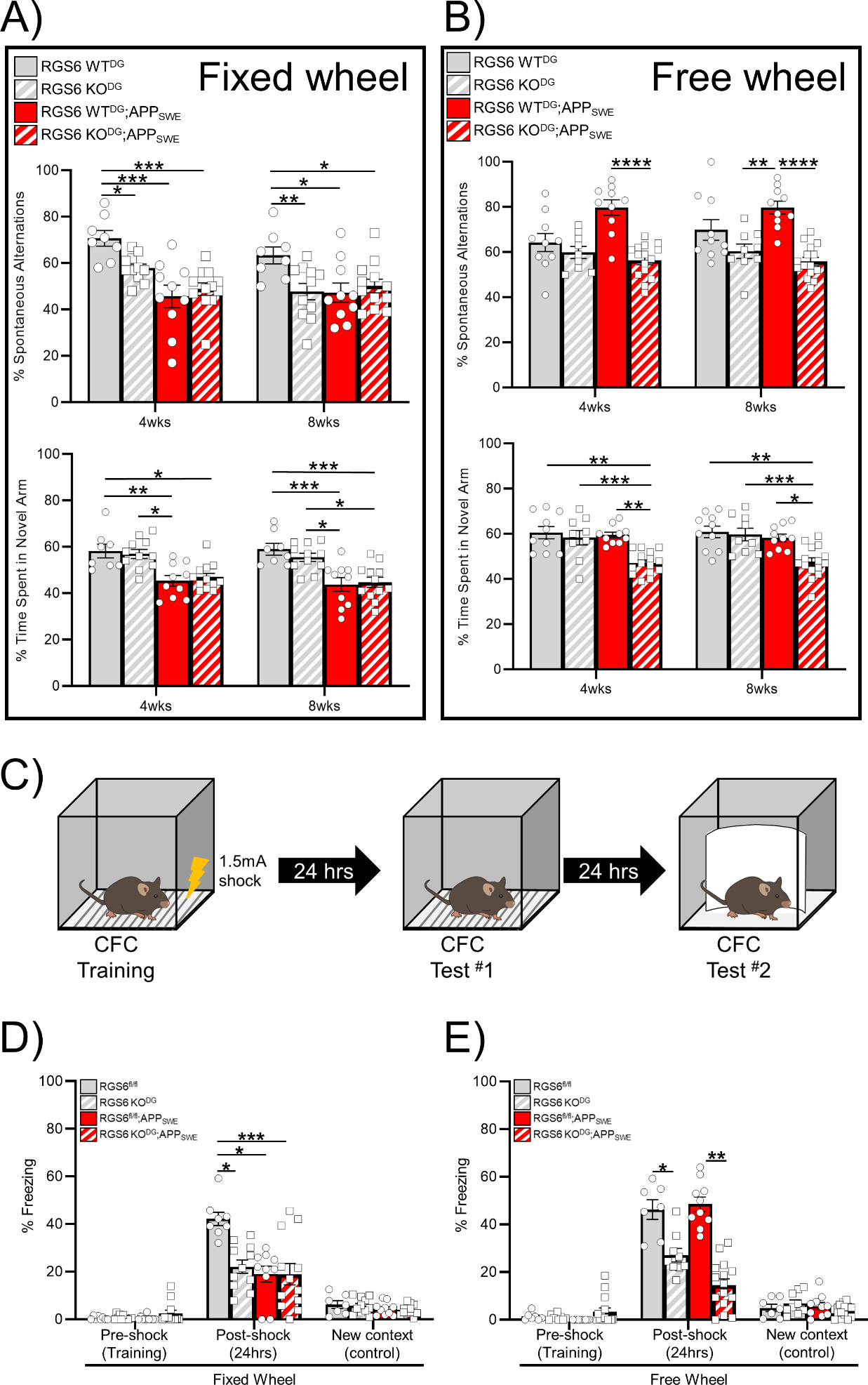
(Related to Fig. 3) *RGS6 in DG NPCs is required for exercise-induced cognitive improvements in APP_SWE_ mice*. **A-B)** % Spontaneous alternations and % time spent in novel arm during 2T Y-maze in mice after 4wks and 8wks on fixed **(A)** or free **(B)** running wheels. **C)** Schematic outlining contextual fear conditioning paradigm. **D-E)** % freezing behavior was recorded before (pre-shock; training) and 24hrs after (post-shock) delivery of a 1.5mA electrical shock to mice. An additional test was run 24hrs from the initial test to ensure freezing behavior was context specific. Percent (%) freezing behavior is shown for mice on fixed wheels **(D)** and free running wheels **(E)**. All post-shock values are significantly different from their respective pre-shock and new context values. Data are expressed as mean ± SEM. Multi-way ANOVA with Tukey’s *post-hoc* analysis was used to analyze the effects of and interactions between genotype, wheel treatment, sex, and time.

**Supplemental Figure 4.**
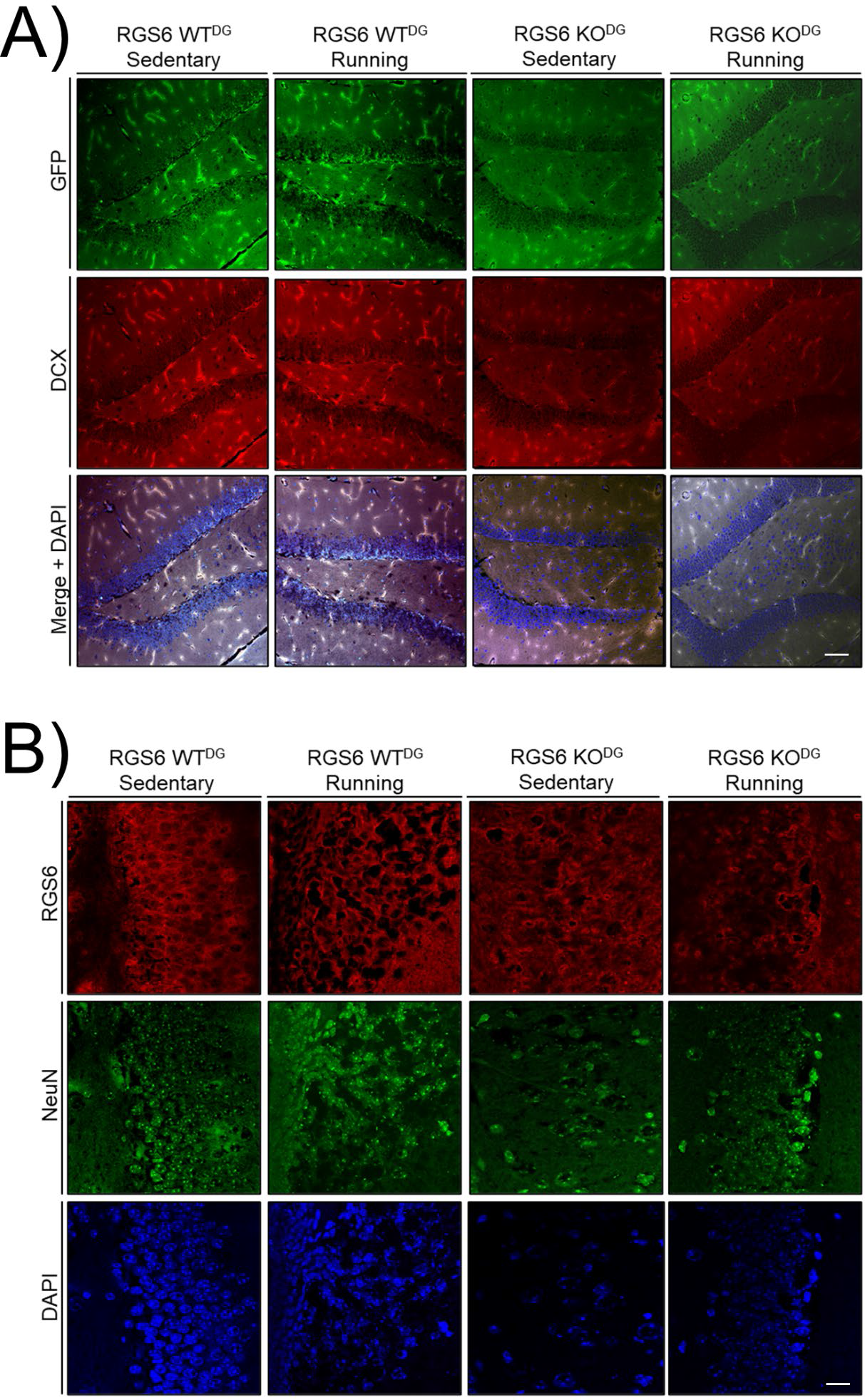
(related to Fig. 4) *RGS6 is necessary for basal and running-induced AHN in APP_SWE_ mice*. **A)** Representative IF images (20x, *N* = 8 RGS6 WT^DG^ animals, *N* = 10 RGS6 KO^DG^ animals) of neurogenesis markers (RV-eGFP or RV-Cre-eGFP, green; doublecortin (DCX), red) and DAPI (blue) in RGS6 WT^DG^ and RGS6 KO^DG^ sedentary and running mice from Figure 3. Scale bar is 25µm. **B)** Representative IF images (40x, *N* = *N* = 8 RGS6 WT^DG^ animals, *N* = 10 RGS6 KO^DG^ animals) analyzing RGS6 (red) co-expression with NeuN (green) and DAPI (blue) in RGS6 WT^DG^ and RGS6 KO^DG^ sedentary and running mice from Figure 3. Scale bar is 40µm.

